# Paraneoplastic renal dysfunction in fly cancer models driven by inflammatory activation of stem cells

**DOI:** 10.1101/2024.03.21.586173

**Authors:** Sze Hang Kwok, Yuejiang Liu, David Bilder, Jung Kim

## Abstract

Tumors can induce systemic disturbances in distant organs, leading to physiological changes that enhance host morbidity. In Drosophila cancer models, tumors have been known for decades to cause hypervolemic ‘bloating’ of the abdominal cavity. Here we use allograft and transgenic tumors to show that hosts display fluid retention associated with autonomously defective secretory capacity of fly renal tubules, which function analogous to those of the human kidney. Excretion from these organs is blocked by abnormal cells that originate from inappropriate activation of normally quiescent renal stem cells (RSCs). Blockage is initiated by IL-6-like oncokines that perturb renal water-transporting cells, and trigger a damage response in RSCs that proceeds pathologically. Thus, a chronic inflammatory state produced by the tumor causes paraneoplastic fluid dysregulation by altering cellular homeostasis of host renal units.

**Significance Statement:** Tumors cause pathophysiological changes to host tissues, including distant organs. Here we use fruit fly cancer models to uncover mechanisms underlying paraneoplastic renal dysfunction. IL-6-like signaling from the tumor induces inflammatory signaling in renal tubule cells. Defects in these cells are sensed by normally quiescent renal stem cells, leading to inappropriate proliferation in a damage-like response. Chronic activation in the tumor context results in physical obstruction of tubule ducts and thus failures in fluid clearance. This fly work can prompt investigation of analogous mechanisms underlying renal dysfunction in cancer patients.

## Introduction

There is growing recognition of the pathological impact of tumors not only on their local microenvironment, but also on the entire body through systemic perturbations. Such ‘paraneoplasias’, which are known to affect muscle, fat, nerves and bone, are major drivers of patient morbidity and mortality but have been much less studied than tumor growth itself (1, 2). Drosophila has recently emerged as a reductionist platform to investigate paraneoplastic mechanisms. Fly cancer models display numerous tumor-host interactions seen in human patients including cachexia, coagulopathy, anorexia and an innate immune response, and many of these are regulated by conserved molecular mediators (3–5). Fly cancer models have also revealed previously unappreciated paraneoplasias that can be cryptically present in mammals as well (6, 7), demonstrating the potential of this system for discovery in cancer biology.

Historically, the first paraneoplasia documented in tumor-bearing flies is a condition in which adult animals present with a dramatically distended abdomen. In a seminal 1969 paper documenting tumor suppressor genes and transplantable malignancies in Drosophila, Gateff termed this phenotype ‘bloating’ (8). Because ‘bloating’ is primarily used by physicians to refer to excess gas, while the fly host abdomen is filled with excess fluid, the phenotype is more accurately described as hypervolemia.

Fluid volume in insects is regulated primarily by renal units called malpighian tubules (MTs), which maintain homeostasis of hemolymph, the circulatory fluid that serves both blood- and lymph-like functions (9). In contrast to the vascular circulation of vertebrates, insects have an open circulatory system in which hemolymph flows freely between internal organs. Water along with metabolic solutes and ions from hemolymph are pumped across the upper region of the MTs into a lumen to create a primary urine. Lower tubules then reabsorb water to condense urine and convey it through a ureter prior to excretion. Transcriptomics reveals that MTs show not only functional but also substantial molecular and cellular analogies to the tubular portion of the kidney, enabling this simple organ to model aspects of human renal biology and pathology (10).

Although described over 50 years ago, only now are mechanisms underlying tumor-induced hypervolemia being investigated. Multiple studies are linking it to defects in the renal system, and highlight the possibility that frequent kidney dysfunction in human cancer patients (11–13) may not be due to chemotherapeutic nephrotoxicity alone. Using transgenic models that transform fly intestinal stem cells, one group has implicated tumor-derived PDGF/VEGF in disrupting function of renal cation-transporting cells, resulting in uric acid accumulation (14). A second group using the same model has identified an antidiuretic hormone produced by the tumor mass that antagonizes fluid absorption by the tubule, and shown that a conserved signaling axis can mediate paraneoplastic renal dysfunction in mammalian cancer (7). Here, using distinct fly tumor models, we identify a new mechanism that disrupts renal tubules via an inflammatory pathway that triggers pathogenic stem cell activity.

## Results

### Abdominal fluid accumulation in fly tumor models is associated with renal secretion defects

To investigate how tumors could disrupt fluid homeostasis, we first reexamined two established adult fly cancer models: an allograft model (**Fig. 1A**) that transplants larval epithelia overexpressing oncogenic *Ras^v12^* and *aPKC^ΔN^* (6) and a new genetic ovarian carcinoma (OC) model in which *Ras^v12^* and *aPKC^ΔN^* are conditionally expressed in the follicular epithelium (4). As previously described and seen also with a genetic intestinal carcinoma (IC) model in which active *yki* is induced in intestinal stem cells (15), this resulted in distended abdomens (**Fig. 1B-F**). Measurements demonstrated a ∼doubling of wet body mass compared to controls; this was supported by a 12 fold-increase in hemolymph volume collected from flies carrying OC (**Fig. 1F, S1A**). The kinetics of fluid accumulation in the models varied, but all tumor-bearing flies showed frank dysregulation before death. Tumors remain predominantly *in situ* and do not contact organs implicated in diuresis (4, 16). These data confirm that fluid imbalance is a syndrome shared in common between tumor-bearing flies. Recognizing that the IC model uses an *esg-GAL4* driver that is expressed in renal cells and an oncogenic responder that is known to induce MT defects (**Fig. 3F, S2A**) (17, 18), we focused on the disc allograft and OC models, which clearly do not drive oncogene expression in renal tissue (**Fig. S1B, C**).

**Figure 1:**
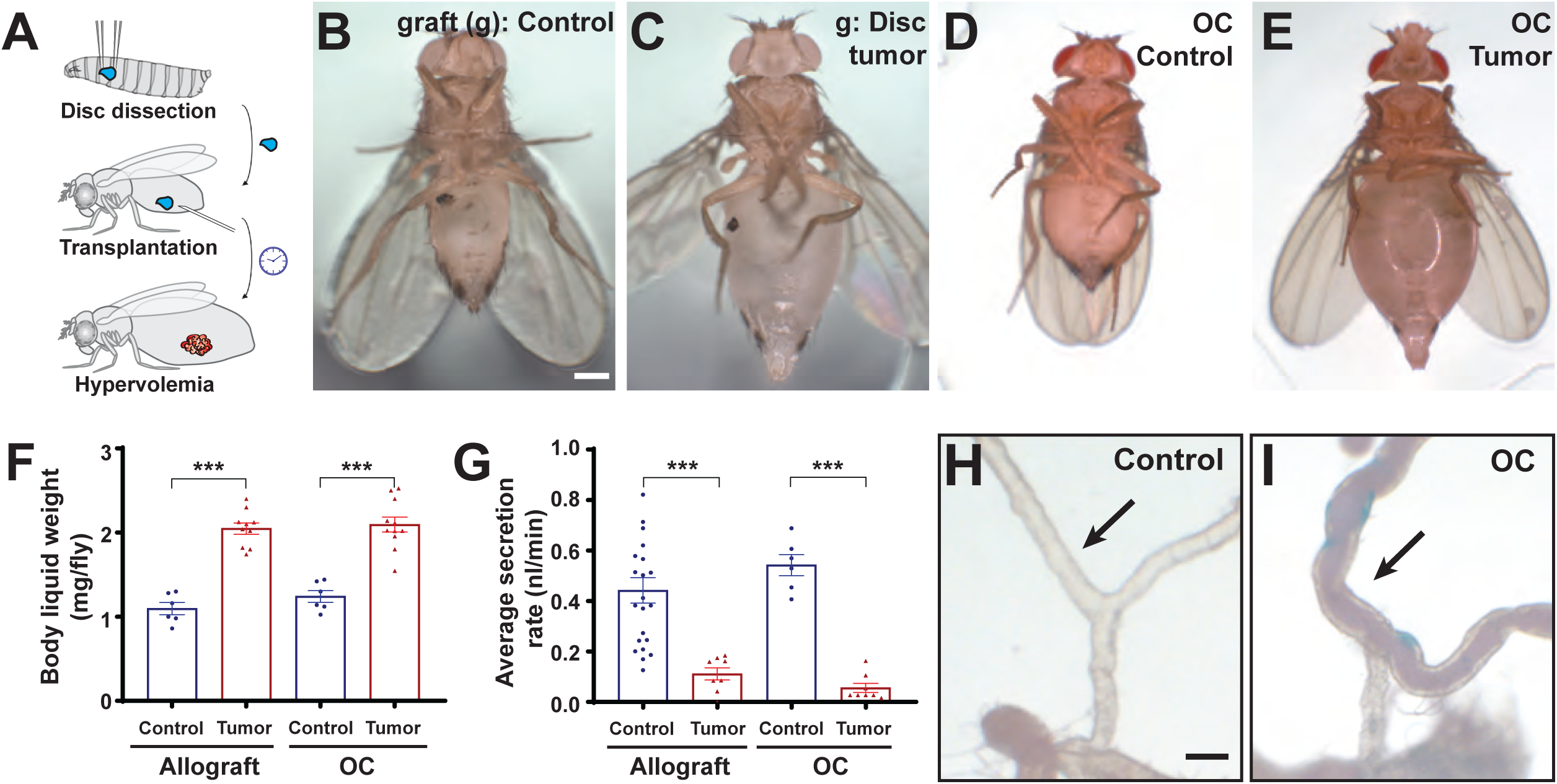
Tumor-induced hypervolemia is associated with defective MT secretion. (**A**) Diagram of allografting into adult flies. (**B-E**) Flies grafted (g, dark spot shows injection site) with a WT disc appear normal (B), while flies grafted with a disc tumor show distended abdomens (C). In genetic OC cancer models, control adults raised at RT appear normal (D), while adults in which tumors are induced by a 29° shift develop abdominal swelling (E). Scale bar, 500 um. (**F**) The phenotype is due to fluid accumulation, as liquid weight is elevated in tumor-bearing flies. (**G**) *Ex vivo* assay demonstrates strongly compromised secretion of MTs isolated from tumor-bearing hosts. (**H, I**) The lower tubules (arrows) from control hosts are clear (H) while those from tumor-bearing hosts are opaque (I). Scale bar, 100 um.

Hypervolemic fluid imbalance could arise from changes in hemolymph composition that lead to osmotic imbalance, as suggested for increased circulating sugars in tumor-bearing fly hosts (15, 19). However, flies exhibiting elevated hemolymph sugar levels due to a high sugar diet or deletion of insulin-like peptides (ILPs) were not hypervolemic (20), suggesting that this mechanism alone cannot account for tumor-induced fluid dysregulation. Moreover, tumor-bearing flies exhibit reduced deposition of urine-containing frass (**Fig. S1E**), a phenotype reminiscent of human patient oliguria that suggests defects in fluid excretion.

To assess whether osmoregulatory hormones were necessary for tumor-induced hypervolemia, we performed Ramsay assays which test the fluid transport capabilities of MTs in *ex vivo* culture (21, 22). Compared to the highly efficient activity seen in wild-type (WT) flies, tumor host MTs showed severely decreased secretion rates (**Fig. 1G, S1D**). Since the Ramsay assay is conducted in media with normal sugar levels and that lacks endocrine signals, the observed defects reflect MT-intrinsic failure rather than the influence of external factors. Therefore, renal tubules of tumor-bearing hosts exhibit organ-autonomous defects.

### Tumor-bearing hosts inappropriately accumulate cells in the tubule lumen

MT functions are governed by two cell types, the principal cells and the stellate cells (9). Principal cells populate the full length of the tubule and provide its major structural basis, whereas stellate cells are located only within the upper tubules (**Fig. 2A, S2A**). Principal cells generate the proton gradient that drives cation and organic solute transport into the MT lumen, whereas stellate cells are the main conduit for water flux.

**Figure 2:**
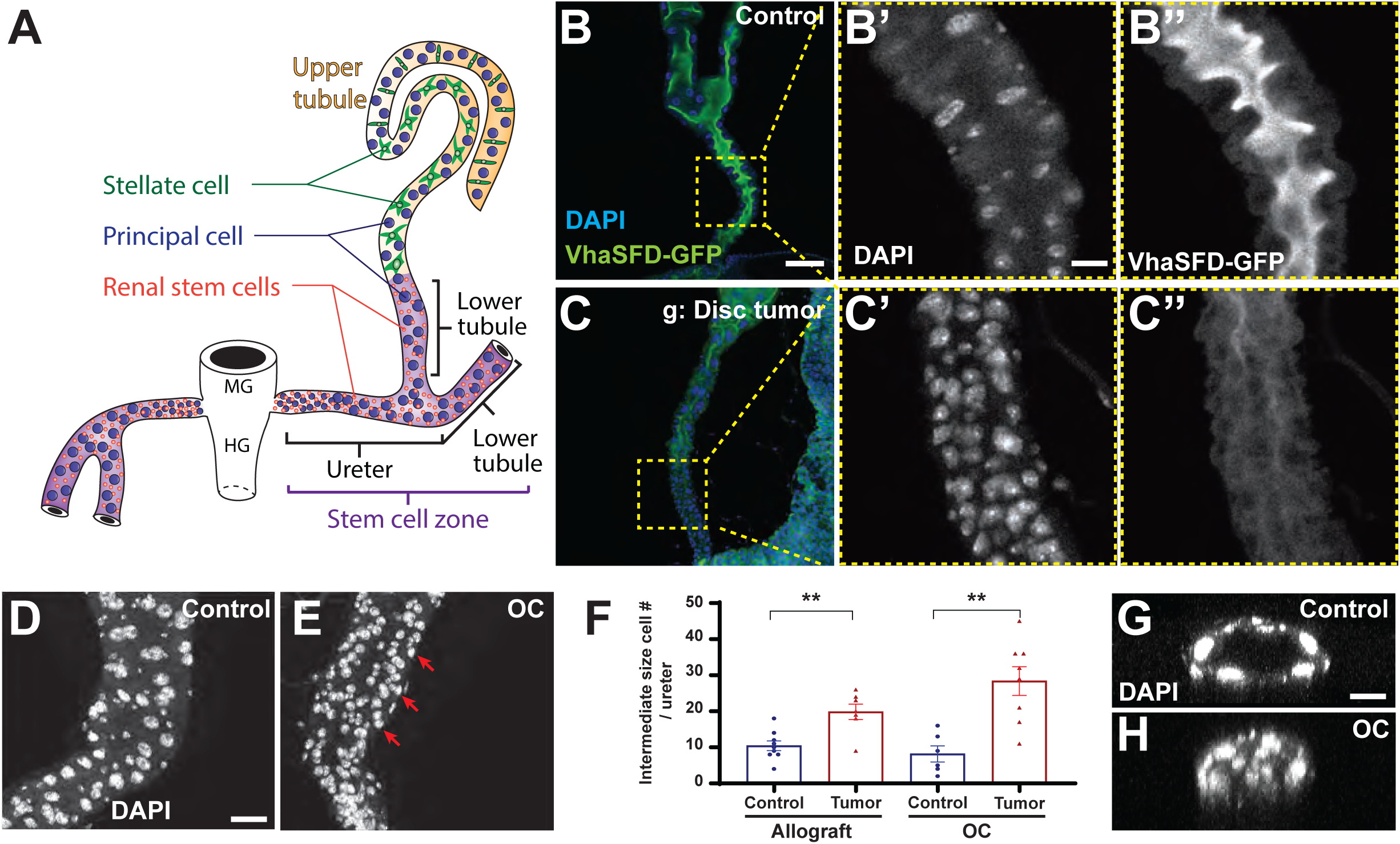
Tumor-bearing hosts accumulate excess cells in the renal tubule lumen. (**A**) Anatomy of MTs, the fly kidney analog (modified with permission from Ref. 21). Tubules absorb fluid from circulation and secrete it into a collecting duct; primary urine moves through the ureter for excretion. (**B-F**) In contrast to the patent lumen of control ureters shown by apical localization of VhaSFD (B, single confocal section), ureters from flies grafted with tumors are occluded by mispolarized cells (C). Control ureters (D, projected confocal stacks) show most nuclei of distinct small and larger sizes, reflecting RSCs and principal cells respectively. Imaging of tumor-bearing flies (E) shows an additional population of intermediate size (red arrows, quantitated in F). Scale bar in B, 100 um. Scale bar in B’ and D, 25 um. (**G, H**) The intermediate-size cells fill the ureter, as seen in cross-sections from control (G) versus tumor-bearing (H) flies. Scale bar, 25 um.

Defective MT secretion in tumor-bearing hosts could be caused by failure of absorption into the tubule or a failure in conveyance through it. Interestingly, brightfield microscopy showed that the lower tubules of tumor-bearing hosts were filled with a cloudy substance reminiscent of condensed urine (**Fig. 1H, I**). This raised the possibility that some tubule transport functions might be intact but excretion via the ureter might be impeded. Gross morphology of the MT was similar in control and tumor-bearing hosts. However, in contrast to the continuous lumen of WT tubules, the lumen of the lower tubules and ureter of tumor-bearing hosts was filled with cells (**Fig. 2B’,C’, E)**. These cells were of a distinct size, smaller than most principal cells from WT tubules (**Fig. 2F, S2B-C**), and their presence was evident as early as 5 days after tumor induction in OC flies. We examined tubules carrying a luminally-localized fluorescently-tagged protein, and found that in tumor-bearing hosts this marker was no longer continuous (**Fig. 2B, C, S2D**). Instead, the ureter lumen contained many misorganized cells, potentially obstructing secretion (**Fig. 2G, H**). We hypothesized that ‘clogging’ of this fluid clearance duct could cause hypervolemia in Drosophila cancer models.

### Tumors trigger abnormal RSC proliferation

What is the source of the luminal cells associated with fluid accumulation? The lower region of MTs, in addition to principal cells, contains small renal stem cells (RSCs) (**Fig. 2A, S2A**). RSCs are normally quiescent but can be activated following injury to replace damaged cells through mitosis, generating new principal cells that can restore tubule function (18, 23). Strikingly, EdU incorporation showed that tumor-bearing host tubules contain proliferating cells, whereas tubules of control flies do not (**Fig. 3A-C**). Because RSCs are the only cell type able to undergo mitosis in adult MTs, we traced cell lineages using G-TRACE (24) driven by *esg-GAL4,* which is expressed in RSCs (**Fig. 3F**) (18). These experiments showed that tumor-induced intermediate size cells were indeed RSC-derived (**Fig. 3D-G**). Interestingly, tumor-bearing hosts did not show obviously greater numbers of RSCs (**Fig. 3F, G**). Instead, lumenal cells expressed the principal cell marker Cut (**Fig. 3H, I)** although levels of other principal cell markers *Alp4-Gal4* and *c42-GAL4* were varied (**Fig. S3A-D**). Based on these phenotypes, we will refer to the intermediate-size lumenal cells as tumor-induced principal cells.

**Figure 3:**
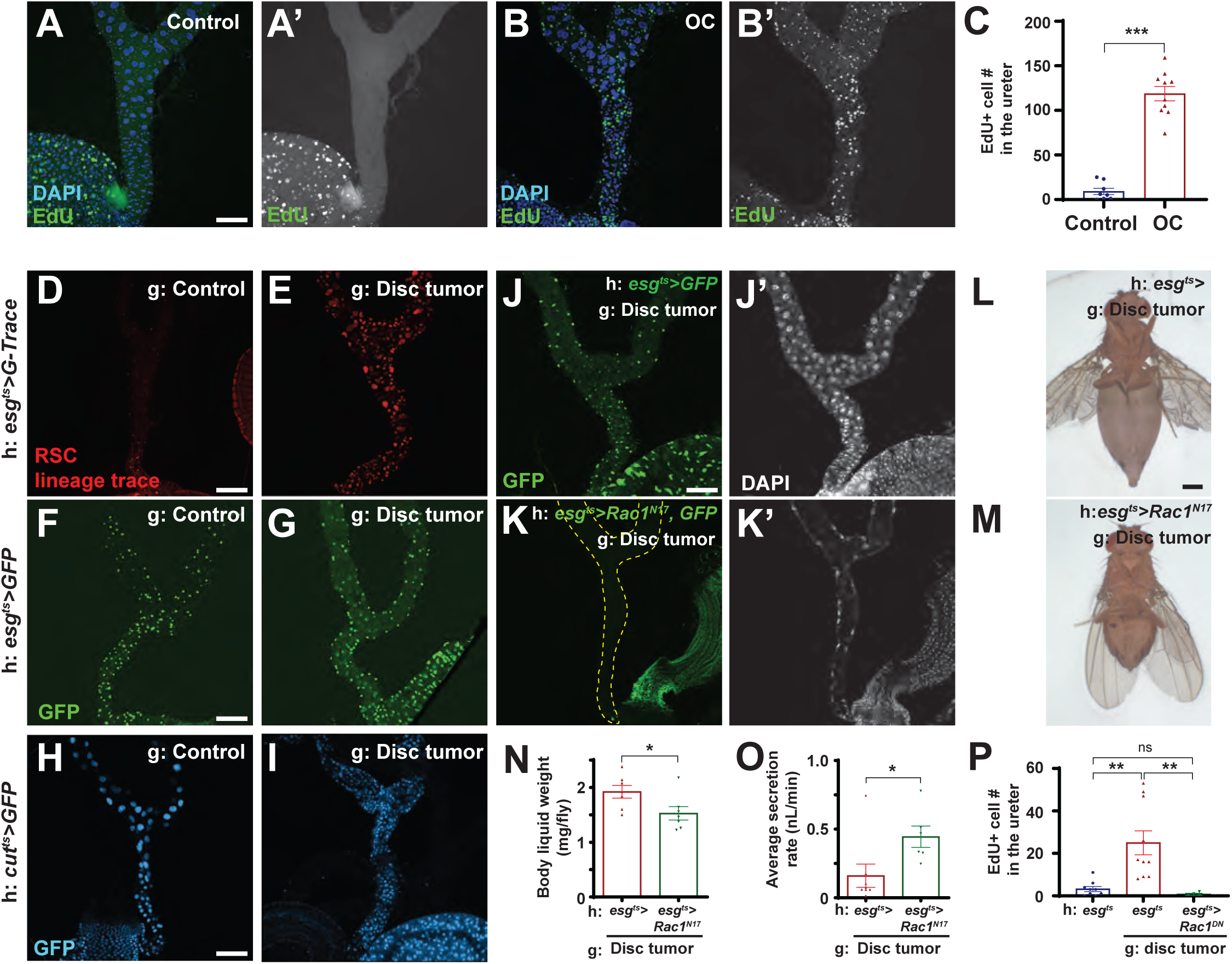
Stem cell-derived progeny are required for fluid imbalance. (**A-C**) EdU incorporation demonstrates cell proliferation in tumor-bearing but not control MTs (A, B, quantitated in C). Scale bar, 100 um. (**D-I**) Ureter-blocking cells are derived from RSCs, as shown by lineage tracing driven by *esg-GAL4* (D, E). However, these cells do not continue expression of the RSC marker *esg-GAL4* (F, G); instead they express the PC marker *Cut-GAL4* (H, I). Scale bar, 100 um. (**J-P**) Presence of RSCs in the MT is blocked by inhibiting Rac1 function during pupation (J, K). Scale bar, 100 um. When grafted with a tumor, these flies fail to develop hypervolemia (L, M, quantitated in N), contain MTs that show rescued fluid transport (O), and lack proliferating cells (P). Scale bar, 500 um.

The identification of tumor-induced principal cells as RSC lineage products provided an opportunity to directly test their role in fluid dysregulation. RSCs migrate from the gut tube to the ureter during pupal stages; this process can be blocked by overexpressing dominant-negative Rac1 in *esg*-positive cells conditionally during pupal stages (17, 23). The animals resulting from this manipulation have grossly normal ISCs in the gut (17) but show RSC-deficient MTs (**Fig. 3J, K**). We therefore transplanted tumors into RSC-deficient hosts and assayed fluid regulation. Indeed, RSC absence prevented proliferation and accumulation of tumor-induced principal cells in host tubules, as predicted by lineage studies above (**Fig. 3J, K, P**). Importantly, RSC absence also rescued tumor-dependent hypervolemia (**Fig. 3L-N**). We confirmed this result by measuring MT secretion rate in the *ex vivo* Ramsay assay, where again robust rescue was seen (**Fig. 3O**). These data support the hypothesis that tubule blockage by RSC-derived cells causes fluid accumulation in tumor-bearing hosts.

### Tumor-induced inflammatory signaling sensed by stellate cells causes fluid retention

JAK/STAT signaling is ectopically activated in several tissues of tumor-bearing hosts, leading to paraneoplasias such as BBB permeabilization and tissue wasting (6, 25). To investigate whether tumors might induce hypervolemia through this inflammatory pathway, we assessed a STAT activity reporter in host MTs. Both control and tumor-bearing hosts showed signal in RSCs. However, only tumor-bearing hosts showed additional expression in stellate cells of the upper tubule as well as cells of the lower tubule and ureter whose size is consistent with tumor-induced principal cells (**Fig. 4A-D**). Interestingly, several changes were evident in the STAT-positive stellate cells of tumor-bearing hosts compared to controls. They lost their distinctive shape and exhibited rounding (**Fig. 4E, F, H, I**). They also reduced expression of differentiation markers such as Tsh and *LKR* (**Fig. 4E-J, S3E**).

**Figure 4:**
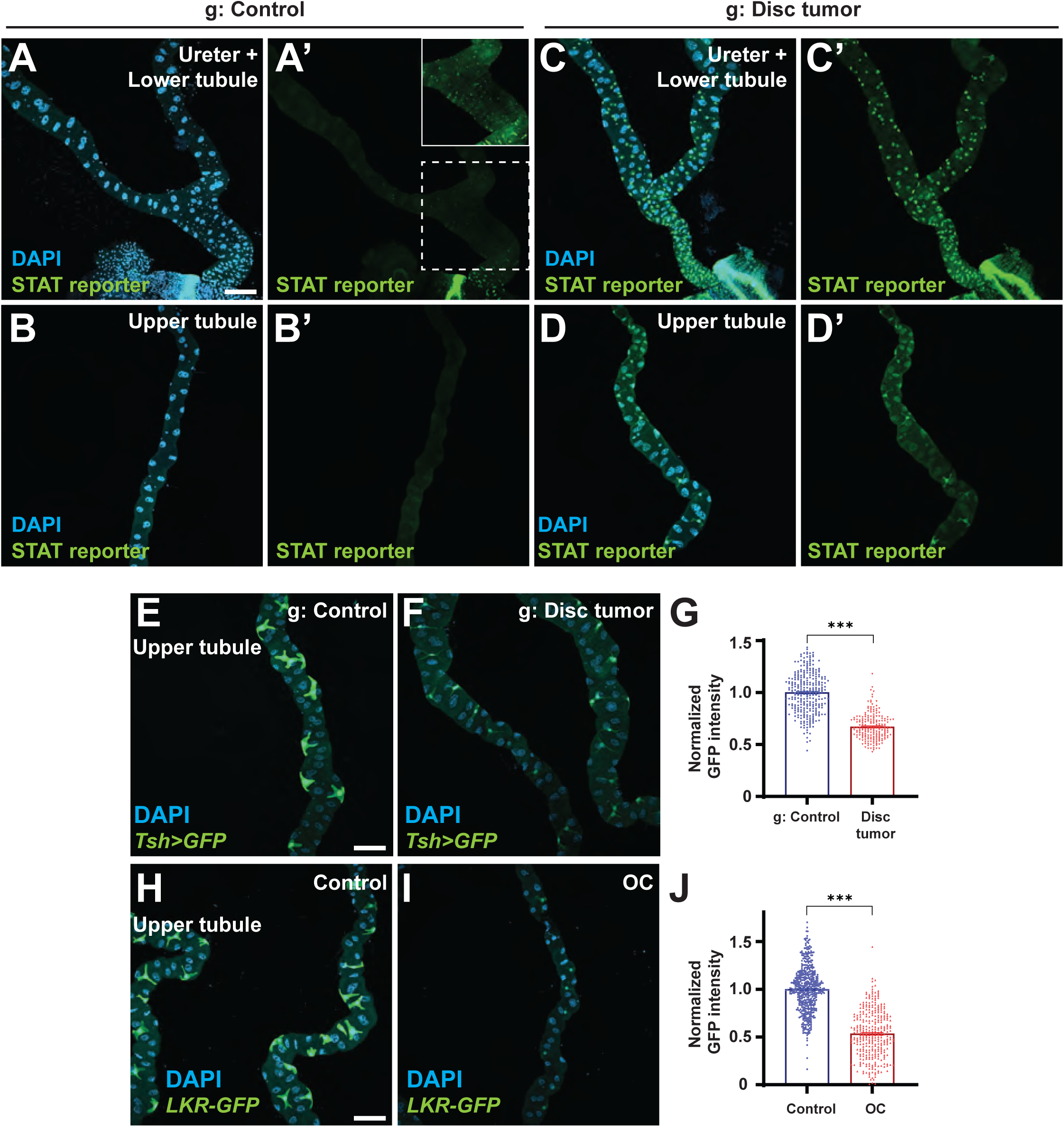
Ectopic STAT signaling in MTs of tumor-bearing hosts. (**A-D**) A reporter for STAT signaling, limited to RSCs (inset, with intensity elevated) in control MTs (A, B), is active in ureter-filling cells (C) as well as stellate cells in tumor-bearing hosts (D). Scale bar, 100 um. (**E-J**) Stellate cells of tumor-bearing flies lose their distinctive shape and reduce expression of markers such as *Tsh>GFP* (E, F, quantitated in G) and *LKR-GFP* (H, I, quantitated in J). Scale bar, 100 um.

We tested the effects of driving constitutively active JAK in different MT cell populations in the absence of tumors. Expression in principal cells or RSCs resulted in no evident defect in fluid balance (**Fig. 5A**). However, expression in stellate cells caused both retention of hemolymph and accumulation of intermediate-sized luminal cells (**Fig. 5A-C**). Morphological and cell fate defects of stellate cells induced by activated JAK were also similar to those seen in tumor-bearing hosts (**Fig. 5D-F**). Thus, STAT signaling in stellate cells can be sufficient to cause fluid dysregulation.

**Figure 5:**
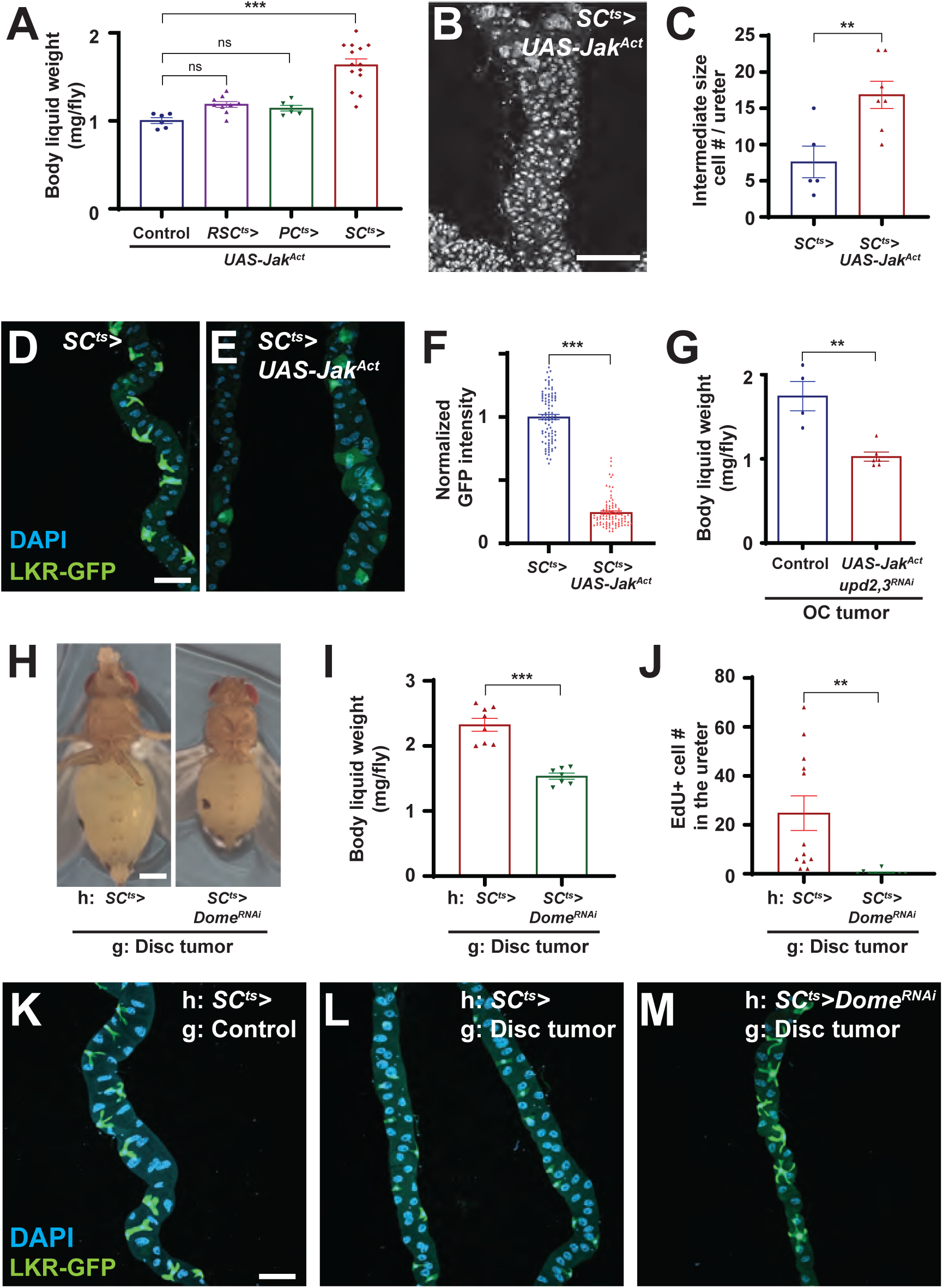
STAT activation in PCs drives renal obstruction. (**A-F**) STAT activation in stellate cells but not principal cells or RSCs is sufficient to cause fluid retention (A), as well as aberrant cells in the ureter (B, C) and defective stellate cells (D, E, quantitated in F). Scale bar, 100 um. (**G**) Hypervolemia is reduced when tumors are depleted of *upd2* and *upd3*, with simultaneous expression of activated JAK to supply tumor-autonomous STAT activation. (**H, I**) Preventing STAT signaling in stellate cells through depletion of Dome reduces hypervolemia of tumor-bearing flies (H), quantitated in (I). Scale bar, 500 um. (**J**) The number of EdU-positive cells in the ureter of tumor-bearing hosts with/without Dome depletion in stellate cells. (**K-M**) Dome depletion also rescues stellate cell morphology and expression of LKR-GFP. Scale bar, 100 um.

To test functional involvement, we allografted disc tumors into hosts in which the STAT pathway receptor Domeless (Dome) was specifically depleted in stellate cells. Strikingly, this led to a rescue of hypervolemia, with wet body weight of tumor-bearing hosts approaching that of control (**Fig. 5H, I)**. It further led to a dramatic decrease of proliferating cells in the ureter (**Fig. 5J**). Finally, it rescued the downregulation of stellate cell fate markers (**Fig. 5K-M, S4A**). By contrast, in tumor-bearing animals whose hypervolemia was rescued by preventing RSC migration and luminal filling, stellate cell fate defects were unchanged (**Fig. S3E**). Together, these data are consiste with a model in which stellate cell dysfunction in response to tumor-driven STAT signaling triggers inappropriate RSC activation, resulting in blocked fluid clearance.

### Upds acts as oncokines on stellate cells that then activate RSCs

Both allograft and ovarian tumors overproduce cytokines of the Unpaired (Upd) family, which are IL-6-like ligands in flies that activate STAT (4, 26, 27). Upd expression can also be induced in WT MTs by damage where it activates RSC proliferation (23). We explored the contribution of Upd sources to fluid imbalance. Allografting disc tumors into *upd2/3-*deficient hosts did not reduce fluid accumulation, suggesting that Upds from host tissues are not required (**Fig. S4B**). Since tumors require autocrine Upds for growth, we restored tumor progression in *upd2/3-* depleted OC cells by co-expressing constitutively active JAK. Flies carrying such tumors, which secrete less Upd into circulation, showed a significant reduction in wet weight compared to flies carrying OC tumors alone (**Fig. 5G**). These data suggest that Upds act as oncokines directly on stellate cells, which then relay another signal to induce RSC proliferation.

How do stellate cells trigger RSC activation? Neither reduction of aquaporin-mediated transport in stellate cells nor stellate cell ablation is sufficient to cause frank hypervolemia (23, 28) (**Fig. S4C**), indicating that the aberrant stellate cells of tumor-bearing hosts display a phenotype other than simple dysfunction. Tubule integrity defects can cause fluid imbalance (29), but cell junctions appeared intact throughout the tubule except at the ureter coincident with cellular disorganization (**Fig. S4D-F**). Pvf1 is natively expressed in stellate cells, and ectopic Pvf1 received in principal cells can result in hypervolemia (14). However, tumor presence failed to enhance stellate cell levels of a Pvf1-GFP reporter (**Fig. S4H, I**). Signaling pathways known to activate RSCs following tubule injury include STAT and MAPK (18, 23). Blocking STAT signaling in RSCs via STAT depletion did not rescue hypervolemia of tumor-bearing hosts (**Fig. S4G**). However, blocking MAPK signaling in RSCs via EGFR depletion resulted in a significant reduction in tumor-induced fluid accumulation (**Fig. S4J**). Tumor-bearing host ureters also displayed elevated levels of dpERK (**Fig. S4K-M**). These data are consistent with a role for MAPK signaling in the process, although how stellate cells stimulate the pathway in RSCs remains to be investigated.

## Discussion

Here we describe a mechanism causing fluid dysregulation in multiple Drosophila cancer models. As with certain human cancer patients (11), the flies present with renal insufficiency and oliguria alongside hypervolemia. The phenotype is a systemic syndrome caused by inflammatory signaling from tumor-derived ligands, which induces a renal damage response and obstruction of fluid clearance ducts. This and contemporaneous work (7, 14) add renal tubules as yet another host organ (alongside fat, muscle, brain and gonads) targeted by direct signaling from fly tumors (3), and allows renal dysfunction to extend the list of paraneoplasias seen in both insect and mammalian tumor-bearing hosts.

Our data including necessity and sufficiency experiments along with RSC depletion implicate the IL-6-like Upd ligands as oncokines initiating signaling defects that ultimately activate RSCs to occlude the renal tubule in both OC and allograft tumor models. This mechanism is distinct from two mechanisms now demonstrated to promote fluid dysregulation in IC tumor models (7, 14). Interestingly, the fly antidiuretic hormone ITP is upregulated in OC as well as IC tumor tissues, and knockdown of ITP in OC diminishes hypervolemia moderately (**Figure S1F**). Thus, an endocrine mechanism, reminiscent of the human Syndrome of Inappropriate Anti-Diuretic Hormone secretion (SIADH), may be a general contributor to paraneoplastic fluid dysregulation. Additionally, ameliorating cachexia-like wasting by knocking down ImpL2 in tumors can partially rescue hypervolemia in the OC and allograft models (**Figure S1F**), as it does in the IC model through unclear mechanisms (15). As now appreciated for cancer cachexia in both flies and humans, different types of tumors or even a single tumor can produce multiple oncokines that trigger the same paraneoplasia through distinct mechanisms. This complexity, which can confound studies in mammalian models, is what the reductionist Drosophila system is well-equipped to untangle.

The ultimate cause of paraneoplastic hypervolemia studied here is inappropriate cellular proliferation, driven by RSCs. The normally quiescent RSCs can be physiologically activated by short-range signals when nearby principal cells in the lower ureter undergo damage (23). A fascinating unanswered question is why RSC activation in response to wounding triggers function-restoring repair, while RSC activation in response to a tumor triggers pathological obstruction.

Since paraneoplasias often reflect corruption of normal interorgan communication, our data suggest that IL-6-like signals from an undefined source may influence osmoregulatory homeostasis, and that stellate cells may be capable of influencing RSC behavior to promote renal tubule maintenance. In addition to physical damage, RSC proliferation may be initiated by stress from MT physiological dysfunction (23, 28, 30); it is possible that this could be triggered by altered stellate cell differentiation seen in tumor-bearing hosts. Like flies, humans can activate a quiescent renal cell population to repair tubules upon injury, and several mouse tumor models that cause renal insufficiency show frank tubular injury alongside endocrine misregulation of physiology (4, 28). Our data provide a basis for investigating non-iatrogenic, paraneoplastic renal dysfunction through disruption of cellular homeostasis in mammalian cancer.

## MATERIALS AND METHODS

### KEY RESOURCE TABLE

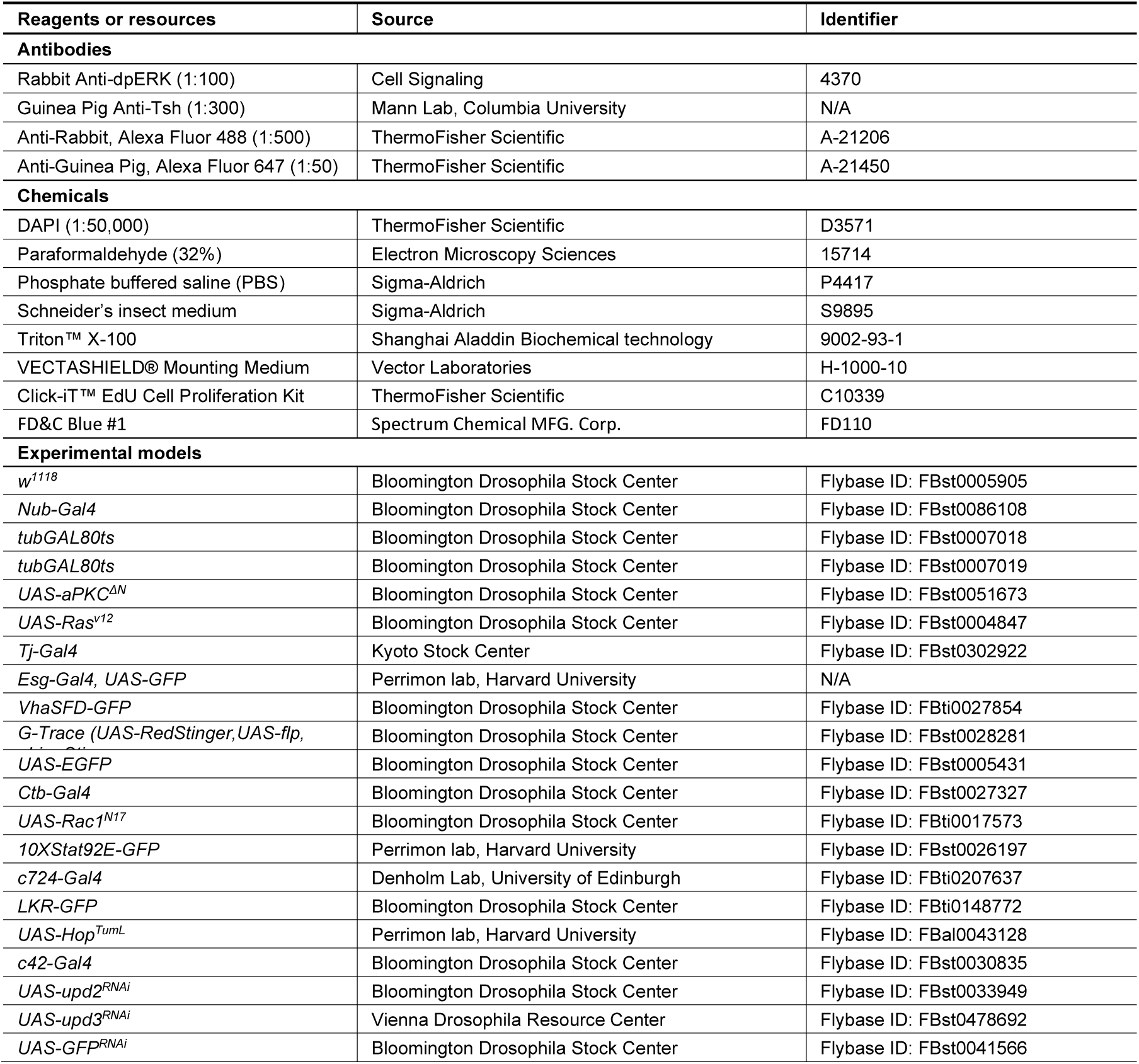
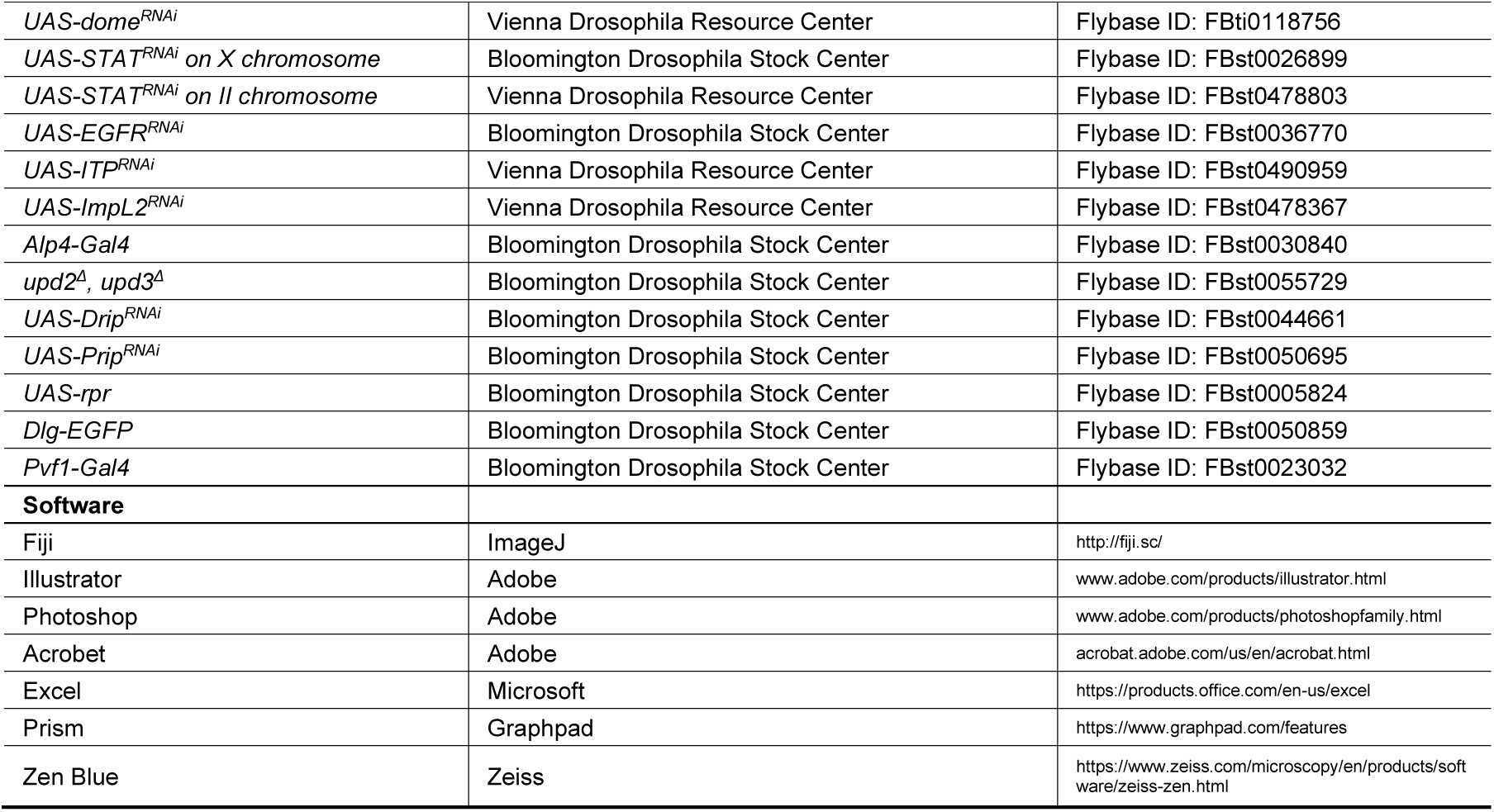

#### Detailed genotypes (related to all figures)

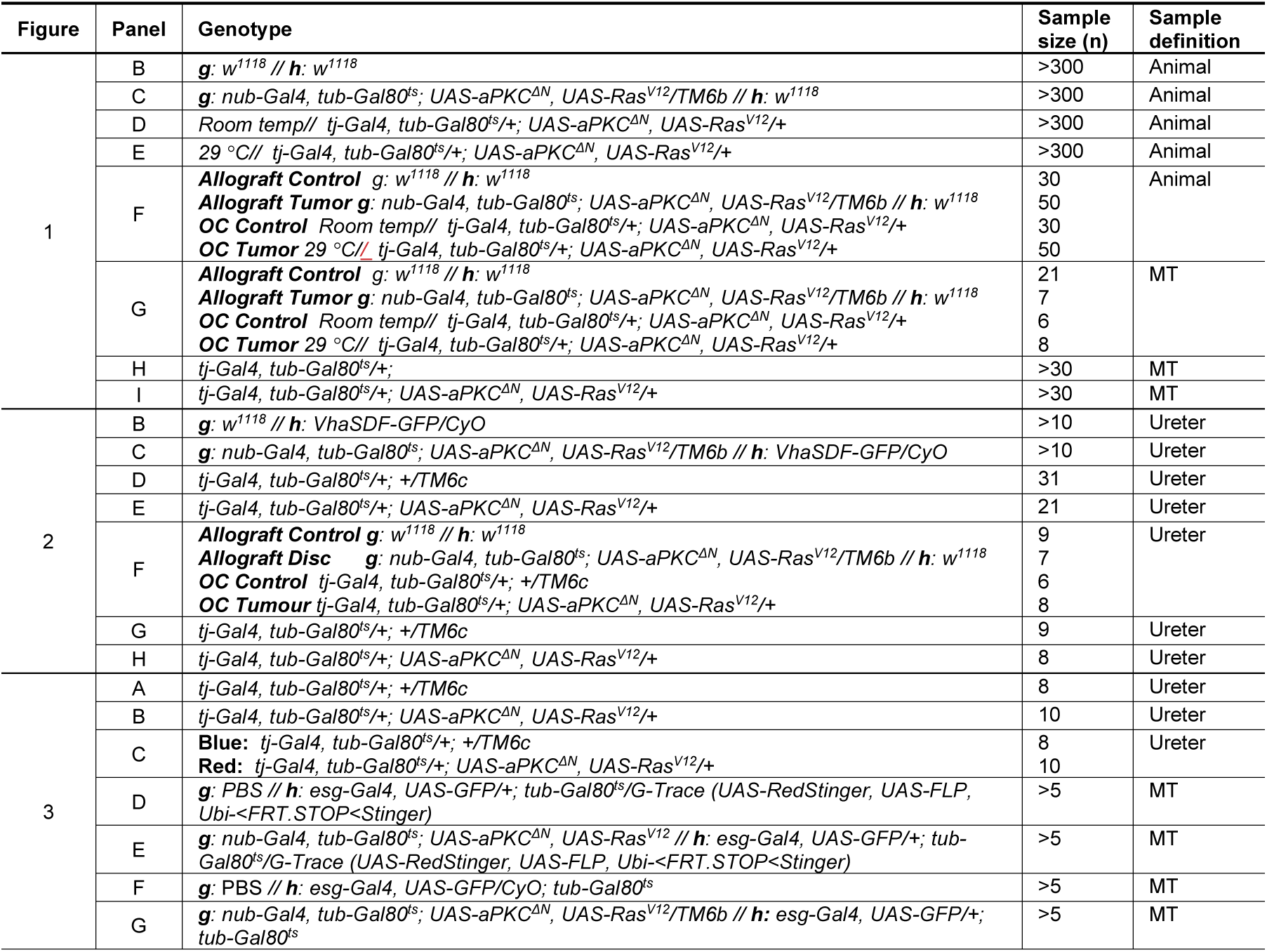
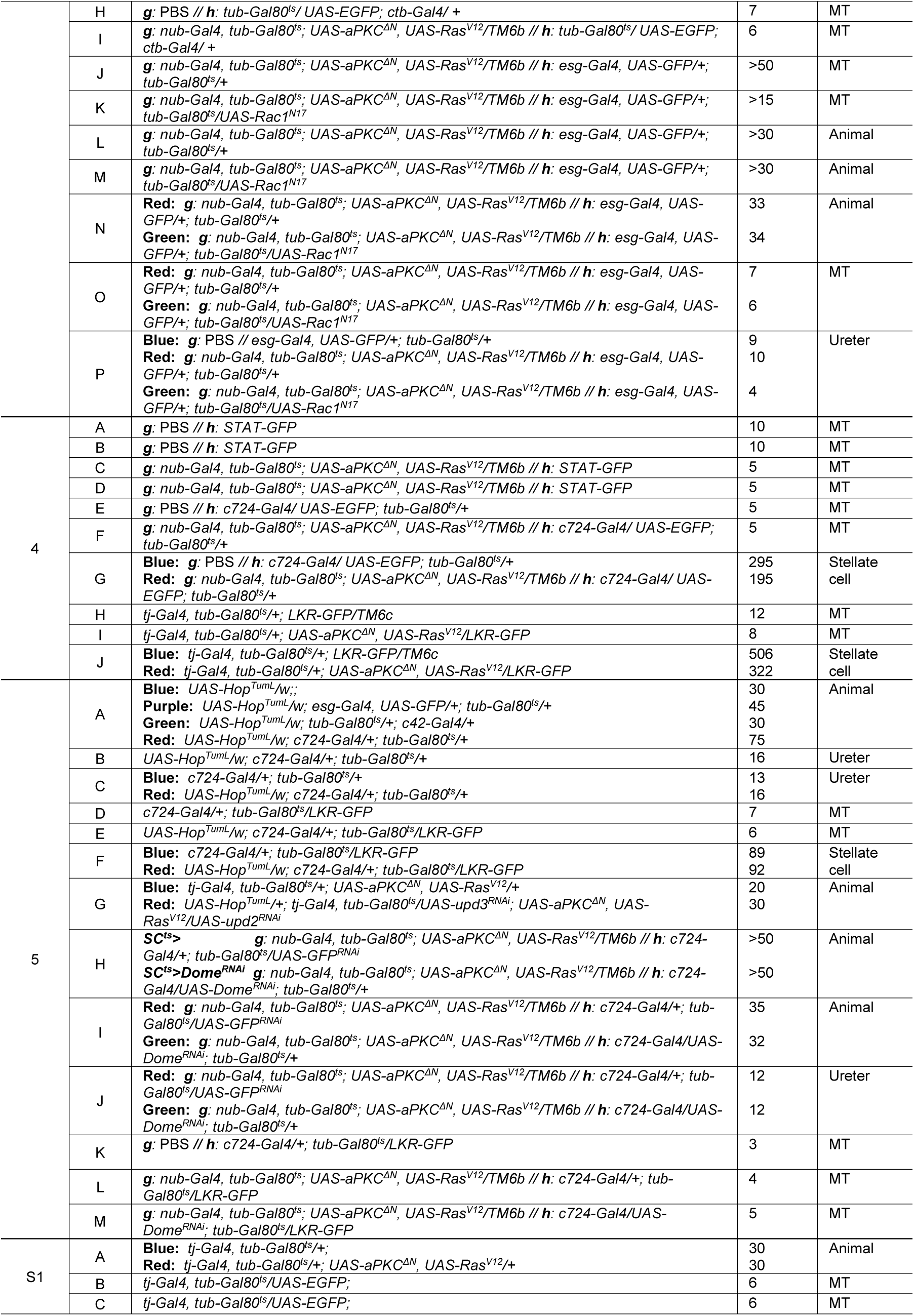
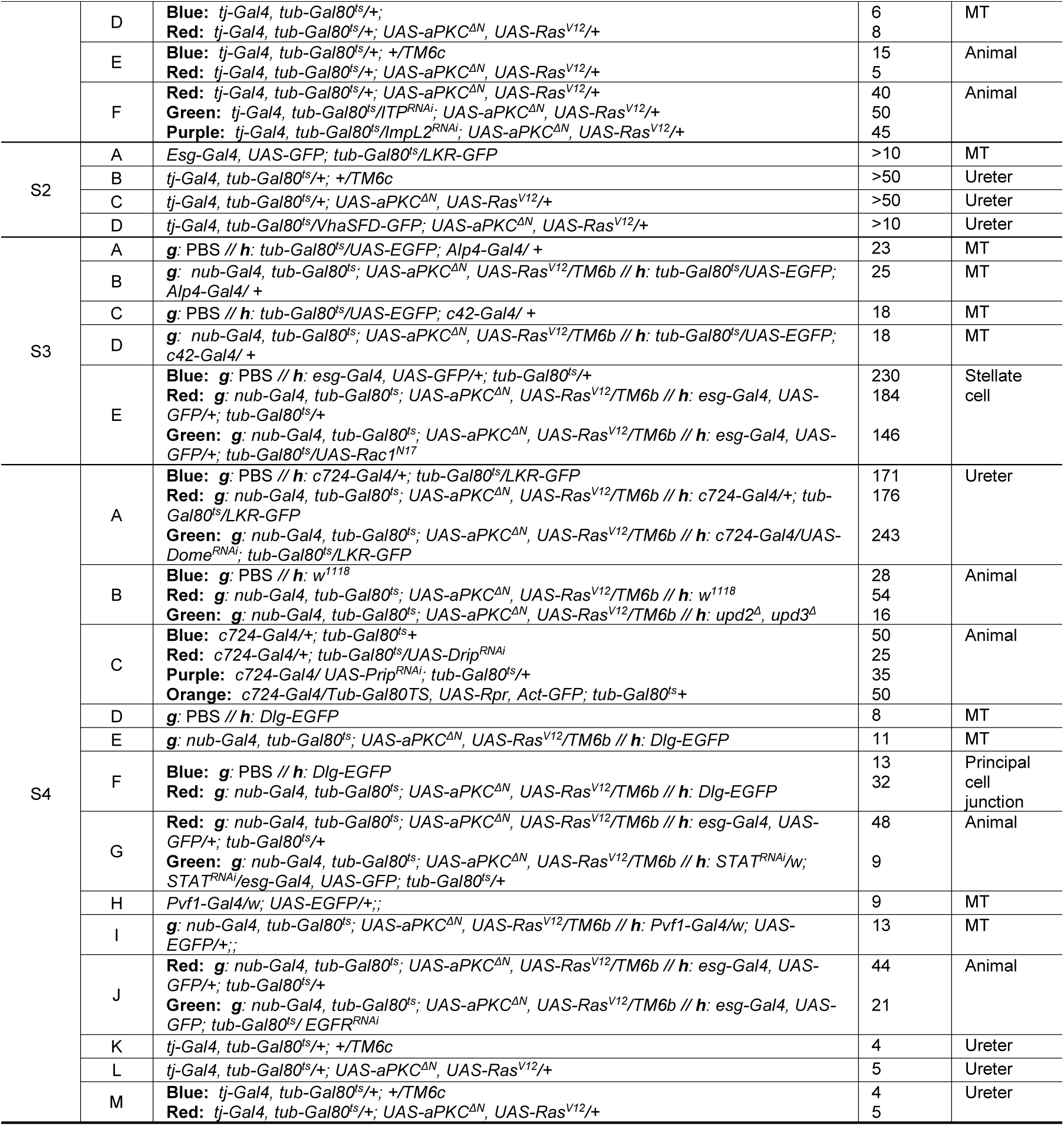

#### Measurement of hemolymph volume and body liquid weight

To collect hemolymph, the abdomen of approximately 10 flies was torn with forceps. These flies were placed in a perforated 0.5-ml Eppendorf tube within a 1.5-ml Eppendorf tube and centrifuged at 2500 rcf for 30 sec. The volume of collected hemolymph was measured with a micropipette P2.5 (Eppendorf) and divided by fly number. To measure body liquid weight, normal weight of 2 - 5 flies was subtracted by dry weight measured after overnight incubation of flies at 65 °C, then divided by fly number.

#### Malpighian tubule secretory assay

The assay was performed as described previously (22). Briefly, a silicone plate was prepared, with a perpendicularly impaled dissection pin adjacent to a ∼10 ul volume indentation. A 1:1 mixture of hemolymph-like saline HL3 (32) and Schneider’s media was loaded in the indentation, and then the wplate was covered by mineral oil. A pair of tubules was dissected by cutting the junction between the gut and the ureter. One tubule was wrapped around a dissection pin and the other tubule was submerged in media. After 15 min (t), the long (a) and the short axis (b) of the ellipsoid-shape media at the tip of the ureter were measured using an ocular micrometer. The secretion rate was estimated by this formula (ellipsoid volume/time): 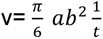.

#### Immunohistochemistry of MT

MTs attached to the gut were fixed in 4% paraformaldehyde for 15 min at room temperature. After washing with PBST (0.3% Triton X in PBS), samples were blocked with PBSTB (2% BSA in PBST) and then incubated with a primary antibody (overnight at 4 °C). After 3 washes with PBST, they were incubated with secondary antibody (for 1 hour at room temperature). After washing, samples were subsequently incubated with DAPI for nuclei staining.

#### Frass assay

Flies were fed with food containing 2.5% w/v blue food dye for 12 h, and then transferred to vials containing normal food for 24h (one fly per vial).

#### Quantification

Images acquired using an LSM900 Inverted confocal microscope (Zeiss) were processed with Zen Blue (Zeiss). Some images were reoriented and non-MT tissues masked for visual clarity. The intensity of the region of interest was quantified using Fiji. For counting the number of tumor-induced principal cells, orthogonal projections of Z-stack images were quantified using Fiji to count DAPI positive staining with the size 20-40 um^2^. Stellate cells were identified manually based on morphology, cell-specific markers, and/or anti-Tsh staining. Images of whole adult flies acquired by Carl Zeiss AxioZoom.V16 were processed with Zen Blue (Zeiss). For MT lumen cross-sections, spacers were used when mounting samples on slides, Z-stack images were processed by Zen Blue (Zeiss) subsequently. Counting of EdU cells was done manually from orthogonal projection of Z-stack images. For the frass assay, excreted dots were counted manually under the dissecting microscope. To measure the intensity of Dlg-GFP, the intensity plot profile of lines (∼10 um) across each bicellular segment of the principal cells was measured using ImageJ.

#### Statistical analysis

The Prism (Graphpad) program was used to run statistical tests. Scatter plots were analyzed using Student’s t-test (comparing 2 groups) or one-way ANOVA with Tukey post-test (comparing more than 3 groups). *P<0.05, **P<0.01, ***P<0.001. Error bar represents mean ± standard error of the mean (sem).

## ACKNOWLEDGMENTS

We thank Iswar Hariharan and Lucy O’Brien for advice and discussion. We acknowledge generous gifts of reagents from Pankaj Kapahi, Barry Denholm, Richard Mann and the community resources provided by TRiP at Harvard Medical School (NIH/NIGMS R01-GM084947), the Bloomington Drosophila Stock Center (NIH P40OD018537) and the Vienna Drosophila Resource Center. This work was supported by NIH grants GM090150 and GM130388 to D.B., and by Seed Fund for Basic Research of the University of Hong Kong to J. K.

