## Supplemental Information for "Paraneoplastic renal dysfunction in fly cancer models driven by inflammatory activation of stem cells"

**This PDF file includes:**

Figures S1 to S4
Supporting Figure Legends

#### Supporting Figure Legends

##### Figure S1

**(A)** Hemolymph volume of OC flies is highly elevated compared to control.
**(B, C)**: T*j-Gal4* lacks expression in MTs; shown are ureter + lower tubules (B) and upper tubules (C). Scale bar, 100 um.
**(D)** Kinetics of fluid transport in Ramsay assays of MTs from control and OC flies.
**(E)** Frass assay shows reduced excretion defects in OC flies compared to control.
**(F)** Depletion of ITP or ImpL2 in OC tumors reduces hypervolemia.

##### Figure S2

**(A)** Cell types in the MT. Scale bar, 100 um. The ureter (A') contains principal cells (PCs) with small nuclei that are comparable in size to the *esg*-expressing RSCs. In the lower tubule (A''), PCs have larger nuclei that lie alongside more RSCs. The upper tubule (A''') lacks RSCs but contains large PCs and the distinctively-shaped stellate cells (SCs), which express *LKR*. Scale bar, 20 um.
**(B, C)** Plot of nuclei sizes in ureters of control (B) and tumor-grafted hosts (C), revealing the latter shows expansion of intermediate-sized cells with nucleus area between 20 – 40 $um^2$.
**(D)** Cells fill disorganized ureters of OC hosts. Scale bar, 100 um.

##### Figure S3

**(A, B)** Abnormal principal-like cells in the ureter of tumor-bearing flies can be Alp4-Gal4 negative. Scale bar, 100 um.
**(C, D)** Abnormal principal-like cells in the ureter of tumor-bearing flies can be c42-Gal4 negative. Scale bar, 100 um.
**(E)** Reduction of Tsh expression in stellate cells of tumor-bearing flies compared to control is not affected by RSC depletion.

##### Figure S4

**(A)** Quantification of Fig. 5K-M.
**(B)** Hypervolemia of tumor-bearing hosts is not rescued when hosts lack *upd2,3*.
**(C)** Depletion of *Drip* or *Prip*, or overexpression of *rpr* is unable to cause hypervolemia.
**(D-F)** Integrity of cell-cell junctions marked by Dlg-GFP appears unchanged in tumor-bearing flies (E) compared to control hosts (D, quantitated in F). Scale bar, 25 um.
**(G)** Hypervolemia of tumor-bearing hosts is not rescued when hosts undergo STAT depletion in RSCs.
**(H, I)** Expression of Pvf1>GFP in stellate cells of tumor-bearing hosts is not elevated compared to control. Scale bar, 100 um.
**(J)** Hypervolemia of tumor-bearing hosts is partially reduced in hosts when EGFR is depleted in RSCs.
**(K-M)** anti-dpERK immunoreactivity at the ureter of OC hosts is elevated compared to control, (quantitated in Q). Scale bar, 100 um.

### Figure S1

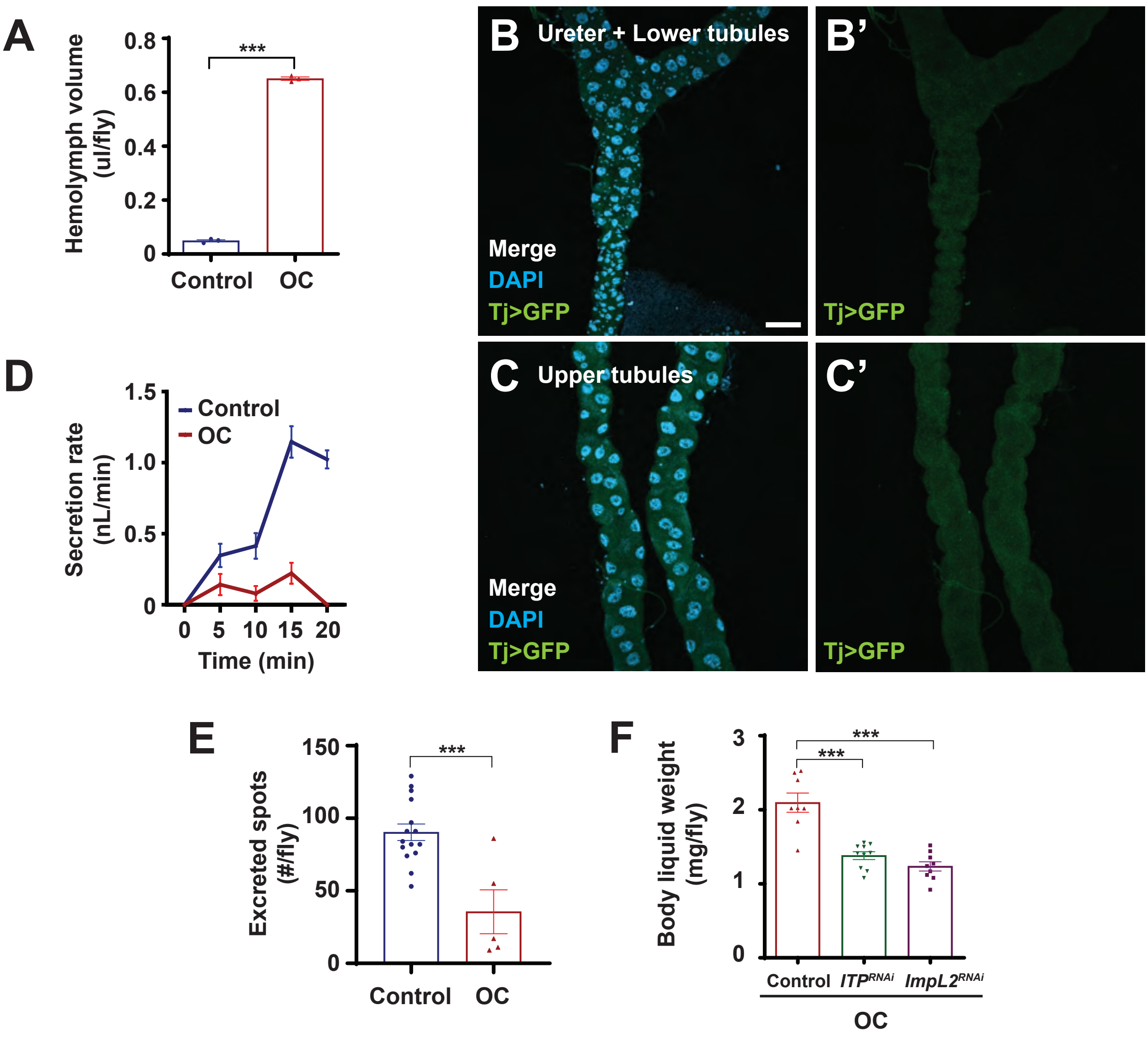
A
Hemolymph volume (ul/fly)
0.8
0.6
0.4
0.2
0
***
Control
OC
B
Ureter + Lower tubules
Merge
DAPI
Tj>GFP
B'
Tj>GFP
C
Upper tubules
Merge
DAPI
Tj>GFP
C'
Tj>GFP
D
Secretion rate (nL/min)
1.5
1.0
0.5
0
Control
OC
0 5 10 15 20
Time (min)
E
Excreted spots (#/fly)
150
100
50
0
***
Control
OC
F
Body liquid weight (mg/fly)
3
2
1
0
***
***
Control
ITPRNAi
ImpL2RNAi
OC

### Figure S2

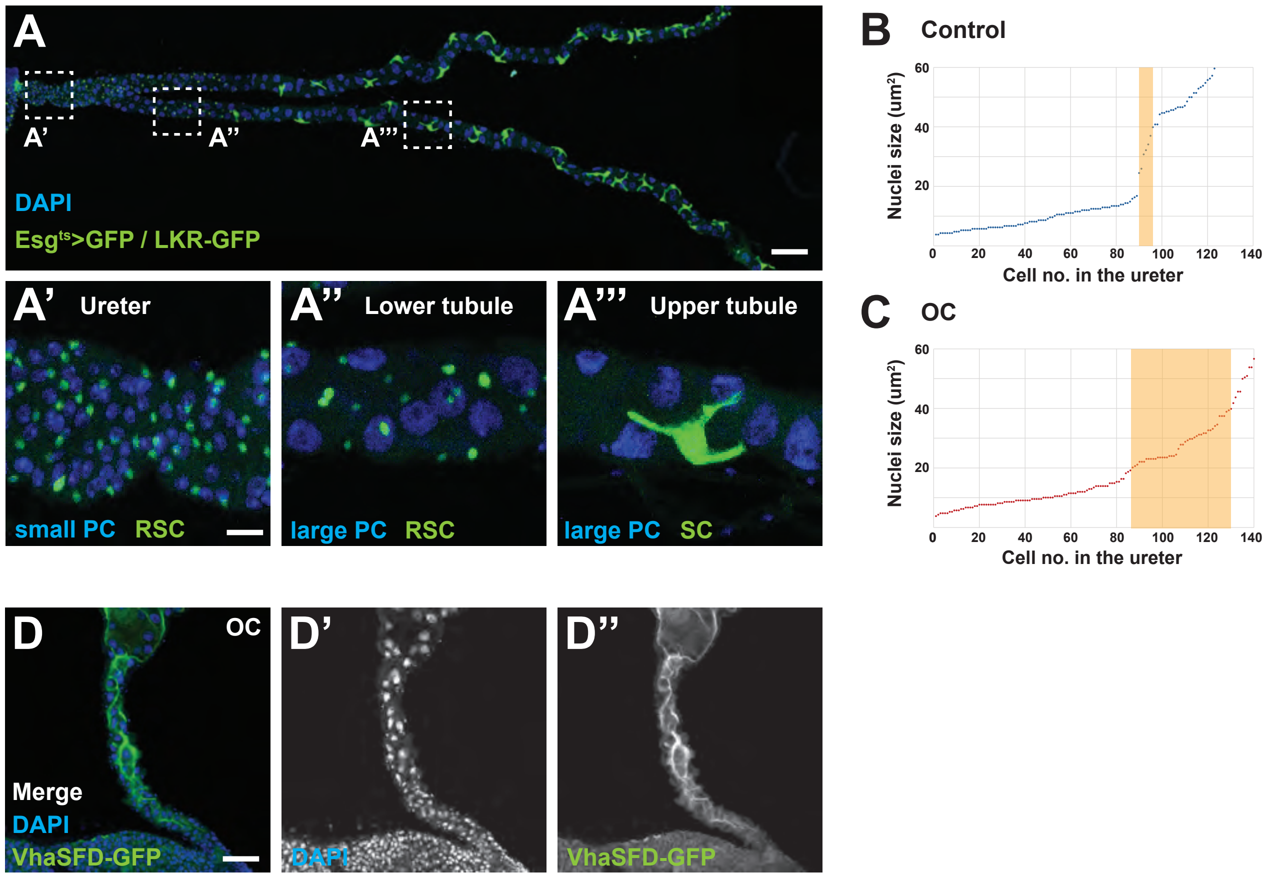

A
A’
A’’
A’’’
DAPI
Esg$^{ts}$>GFP / LKR-GFP
A’ Ureter
small PC RSC
A’’ Lower tubule
large PC RSC
A’’’ Upper tubule
large PC SC
B Control
Nuclei size (um$^2$)
Cell no. in the ureter
C OC
Nuclei size (um$^2$)
Cell no. in the ureter
D OC
Merge
DAPI
VhaSFD-GFP
D’
DAPI
D’’
VhaSFD-GFP

### Figure S3

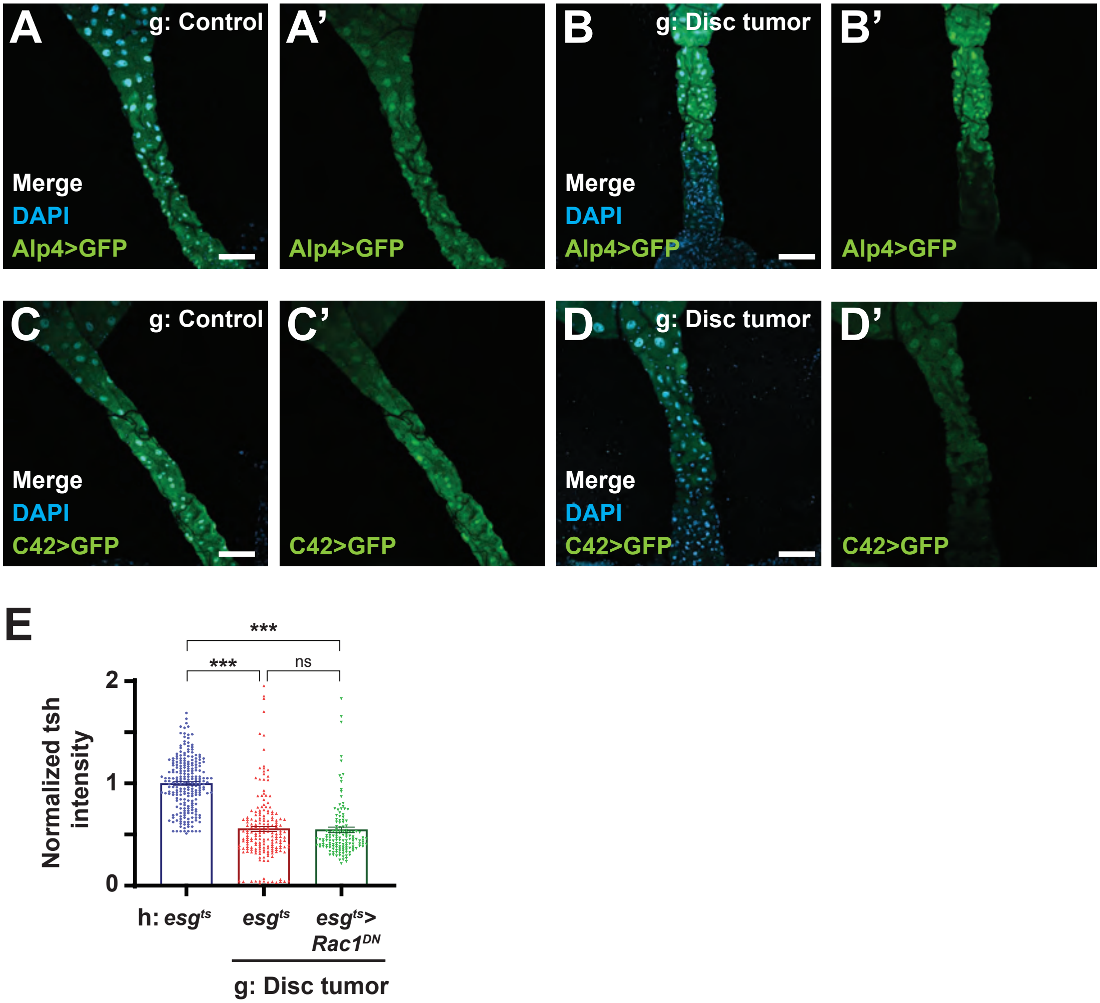
A
g: Control
Merge
DAPI
Alp4>GFP
A'
Alp4>GFP
B
g: Disc tumor
Merge
DAPI
Alp4>GFP
B'
Alp4>GFP
C
g: Control
Merge
DAPI
C42>GFP
C'
C42>GFP
D
g: Disc tumor
Merge
DAPI
C42>GFP
D'
C42>GFP
E
***
***
ns
Normalized tsh intensity
2
1
0
h: esgts
esgts
esgts> Rac1DN
g: Disc tumor

### Figure S4

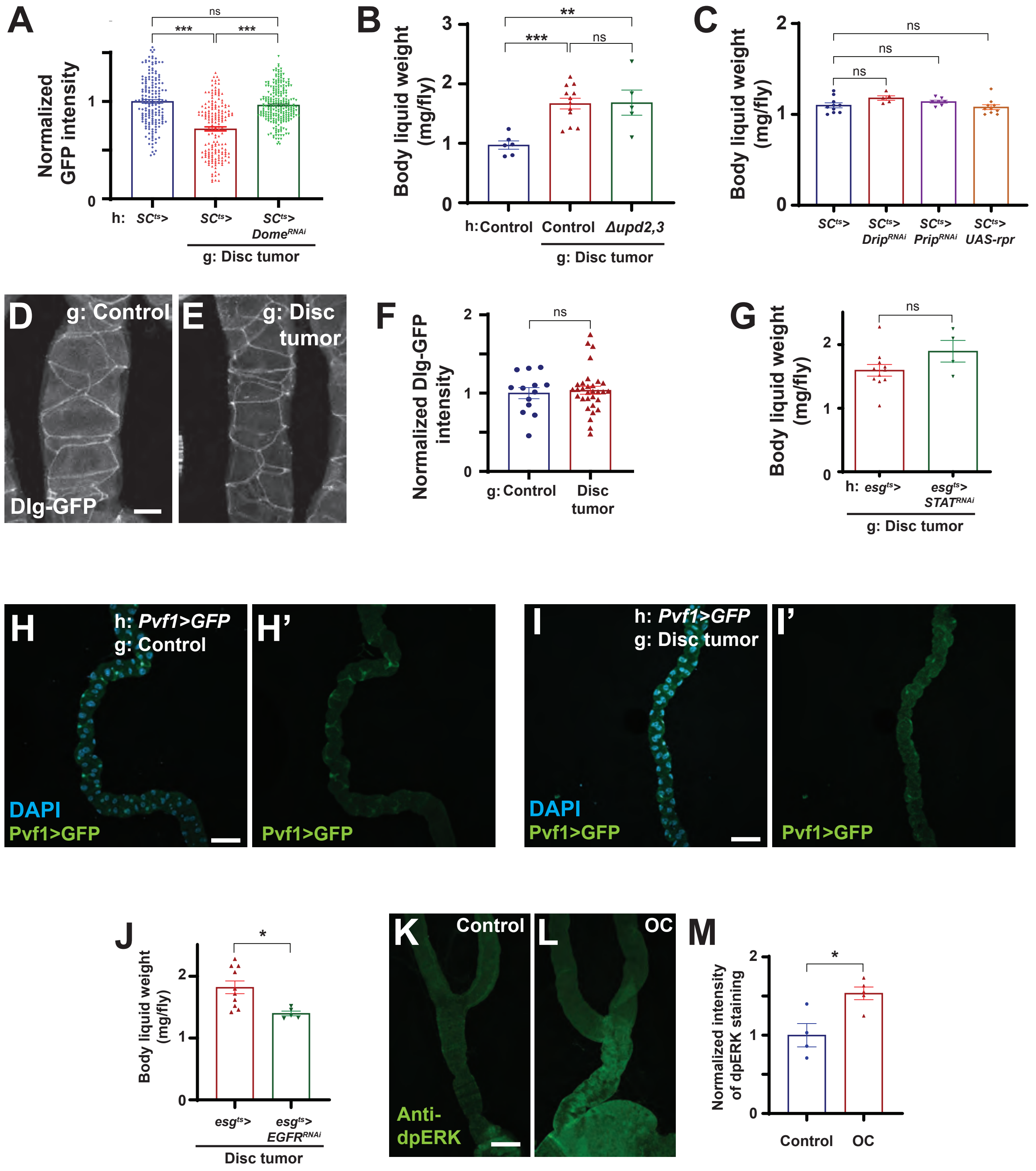

A
Normalized GFP intensity
ns
***
***
h: SC^ts>
SC^ts>
SC^ts> Dome^RNAi
g: Disc tumor
B
Body liquid weight (mg/fly)
**
***
ns
h: Control
Control
Δupd2,3
g: Disc tumor
C
Body liquid weight (mg/fly)
ns
ns
ns
SC^ts>
SC^ts> Drip^RNAi
SC^ts> Prip^RNAi
SC^ts> UAS-rpr
D
g: Control
Dlg-GFP
E
g: Disc tumor
F
Normalized Dlg-GFP intensity
ns
g: Control
Disc tumor
G
Body liquid weight (mg/fly)
ns
h: esg^ts>
esg^ts> STAT^RNAi
g: Disc tumor
H
h: Pvf1>GFP
g: Control
DAPI
Pvf1>GFP
H'
Pvf1>GFP
I
h: Pvf1>GFP
g: Disc tumor
DAPI
Pvf1>GFP
I'
Pvf1>GFP
J
Body liquid weight (mg/fly)
*
esg^ts>
esg^ts> EGFR^RNAi
Disc tumor
K
Control
Anti-dpERK
L
OC
M
Normalized intensity of dpERK staining
*
Control
OC
